# EBSeq: improving mixing computations for multi-group differential expression analysis

**DOI:** 10.1101/2020.06.19.162180

**Authors:** Xiuyu Ma, Christina Kendziorski, Michael A. Newton

**Affiliations:** Department of Statistics, University of Wisconsin Madison, 1300 University Ave, 53706 WI, USA; Department of Biostatistics and Medical Informatics, University of Wisconsin Madison, 425g Henry Mall, 53706 WI, USA; Department of Biostatistics and Medical Informatics, and Department of Statistics, University of Wisconsin Madison, 1300 University Ave, 53706 WI, USA

**Keywords:** Empirical Bayes, hierarchical model, Bayes factor, pruning, local false discovery rate

## Abstract

**EBSeq** is a Bioconductor package designed to calculate empirical-Bayesian inference summaries from sequence-based gene-expression (RNA-Seq) data. It produces gene or isoform-specific scores that measure various patterns of differential expression among a set of sample groups, and is most commonly deployed to measure differential expression between two groups. Its use of local posterior probabilities from a fitted mixture model provides the data analyst a direct way to score the false discovery rate of any reported list of genes, and it is one of the only tools that can address local false discovery rates when analyzing multiple sample groups. Contemporary applications have increasing numbers of sample groups, and the algorithms deployed in **EBSeq** are neither space nor time efficient in this important case. We describe a version update utilizing code improvements and novel pruning and clustering algorithms in order to reduce the complexity of mixture computations. The algorithms are supported by a theoretical analysis and tested empirically on a variety of benchmark and synthetic data sets.

## Introduction

^1^ Since their introduction over twenty years ago, technologies to measure genome-wide gene expression have revolutionized science and medicine. The resulting data have had a major impact on statistical sciences as well, by introducing challenges arising from “small n, large p” datasets. One of the central statistical challenges has been the differential expression problem: namely, how do we identify genes whose expression levels vary significantly across biological conditions? Dozens of methods have been developed toward this end and a few have endured. For RNA-Seq data, the empirical Bayesian hierarchical modeling approach, encoded in R package **EBSeq** has a number of advantages owing to how it captures variation characteristics of genes and isoforms and how it scores differential expression over two or more conditions (Leng, Dawson, Thomson, Rissman, Smits, Haag, Gould, Stewart, and Kendziorski 2013). It has proven useful in hundreds of studies including studies of development (Louro, Marques, Manchado, Power, and Campinho 2020; Sanders and Cartwright 2015; Yoon, Ringeling, Vissers, Jacob, Pokrass, Jimenez-Cyrus, Su, Kim, Zhu, Zheng *et al.* 2017; Sabbagh, Heng, Luo, Castanon, Nery, Rattner, Goff, Ecker, and Nathans 2018), viral transcription (Newhouse, Hofmeister, and Balakrishnan 2017; O’Grady, Baddoo, and Flemington 2017; Baños-Lara, Zabaleta, Garai, Baddoo, and Guerrero-Plata 2018; Zhang, Zeng, Zhou, Irving, Li, Shi, and Wang 2017), and cancer (Lee, Ahn, Jeong, Pak, Moon, and Kim 2020; Song, Tang, Li, Liu, and Zhou 2018; Son, Jeong, Park, No, Lee, Lee, and Chung 2017; Yang, Lv, Lv, Liu, Dong, Yao, Dai, Zhang, and Wang 2016). **EBSeq** has been deployed in Bioconductor since 2013 (Leng and Kendziorski 2019; Huber, Carey, Gentleman, Anders, Carlson, Carvalho, Bravo, Davis, Gatto, Girke, Gottardo, Hahne, Hansen, Irizarry, Lawrence, Love, MacDonald, Obenchain, Ole’s, Pag’es, Reyes, Shannon, Smyth, Tenenbaum, Waldron, and Morgan 2015).

The most common use of **EBSeq** is to score differential expression between two biological conditions. The package’s multi-condition feature is less frequently deployed; but recently it has been recognized that **EBSeq** multi-condition scores are uniquely suited to characterizing multiple cellular subtypes from the growing body of single-cell RNA-seq data (Ma, Korthauer, Kendziorski, and Newton 2019). The work reported here is motivated by the need to improve computational performance of the multi-condition calculations within the original system, say **EBSeq.v1**, which at the time of writing is version 1.26.0 at Bioconductor. In addition to some basic code improvements, we deploy in **EBSeq.v2** a new algorithmic approach to determining multi-condition differential expression scores.

With samples from *K* biological conditions, **EBSeq.v1** calculates posterior probability scores for various patterns of differential expression among these conditions. Typically the null pattern, in which expected expression is the same across all groups, receives the highest score, on the average over genes; the software can consider many patterns. A computational bottleneck arises if we ask the code to consider all possible patterns, which are equal in number to the Bell number, *B*_*K*_, of partitions of *K* objects (Gardner 1978). For even moderate *K*, the memory and time costs of **EBSeq.v1** become excessive. Section 2 describes an alternative pruning/clustering algorithm which leverages the finding that many of the differential expression patterns will have small mixing rates; and we can know this without fitting the full model. By identifying patterns that are probably inconsequential to the final inferences, we remove them from other compute-intensive parts of the code and improve the overall operating characteristics, as demonstrated in a battery of tests in Section 3.

**EBSeq.v2** improves the performance of **EBSeq.v1**. In addition to improved handling of group partitions indicated above and discussed in Section 2, the core code is converted to C++ and adopts open-source, peer-reviewed, and fast libraries **Eigen** and **Boost** for internals (Guennebaud, Jacob *et al.* 2010; Boost 2015). We also modify the EM algorithm by changing how the hyperparameters are recomputed in each cycle. We substitute the Nelder-Mead optimization (`optim` from package **stats**) with a single gradient step within each EM update. The overall effect of these changes is to dramatically improve the performance of **EBSeq** in the multi-group setting. Section 3 presents a variety of simulated and empirical examples and a discussion follows in Section 4.

## METHODS

### The statistical problem

For each inference unit in the system we have real-valued measurements *X* = {*X*_*i*_}, for a sample index *i* = 1, 2,…, *n*, as well as discrete sample labels, say *y*_*i*_, taking values in a label set {1, 2,…, *K*}. The labels refer to different sampling groups, or conditions, that underlie the measurements, and we are especially interested in the case when *K* exceeds 2. In applications of interest we expect *K* to be small compared to *n*, perhaps in the tens when *n* is in the hundreds to thousands. Statistical inference is focused on evaluating hypotheses about the unknown mean values *µ*_*k*_ = *E*(*X*_*i*_|*y*_*i*_ = *k*). For example, in the case of *K* = 4 groups, we have *B*_4_ = 15 different patterns of equality and inequality among the group means, a few of which are:

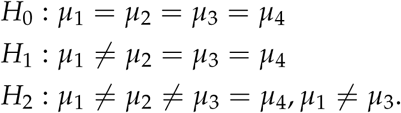

The set of partitions of {1, 2,, *K*} is in one-one correspondence with the set of such hypotheses regarding equalities and inequalities among the means. More specifically, for a partition *π* = {*b*} composed of mutually disjoint blocks *b* ⊂ {1, 2,, *K*}, we say that mean vector *µ* = (*µ*_*k*_) satisfies pattern *π* if *µ*_*j*_ = *µ*_*k*_ whenever *j, k* ∈ *b* for some *b* ∈ *π* and also if *µ*_*j*_ ≠ *µ*_*k*_ whenever *j* and *k* are in different blocks. Thus, for example, *H*_0_ above corresponds to *π* = {{1, 2, 3, 4}}, and *H*_1_ corresponds to *π* = {{1}, {2, 3, 4}}. In the remainder, we use notation *M*_*π*_ to denote the the constraint (i.e., hypothesis) on the mean vector associated with partition *π*:

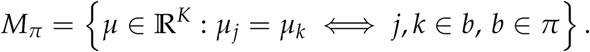

Notice that any mean vector *µ* is an element of exactly one of these subsets, and, considering all partitions,

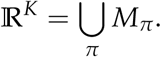

The use of partition/pattern structures for expected values appears in many statistical problems. They are a central feature of Dirichlet-process mixture computations (MacEachern 1998), various Bayesian clustering algorithms (Quintana and Iglesias 2003; Heller and Ghahramani 2005), as well as empirical Bayesian methodologies for genomic applications, such as the **EBSeq** tool introduced earlier, and a similar tool for microarray-based data, **EBarrays** (Kendziorski, Newton, Lan, and Gould 2003; Yuan, Newton, Sarkar, and Kendziorski 2019).

In empirical Bayesian applications, there are very many inference units (genes), and for each sample *i* there is a measurement on every unit. The methodology entails both discrete and continuous mixing over the parameter space in order to deliver posterior probability scores for each unit: *p*_local,*π*_ = *P*(*M*_*π*_|*X, y*). We use the subscript ‘local’ to remind us that the the probability is local to the particular unit (e.g., gene), in the same way that the local false discovery rate is local to each testing unit in large-scale inference (Efron 2010).

The discrete mixing further involves probabilities estimated for the whole system, *p*_global,*π*_. By the use of maximum likelihood estimation, the fitted probabilities satisfy:

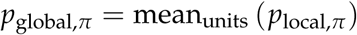

The conditional independence assumptions behind **EBSeq** induce a product-partition form on *p*_local,*π*_: with {*X*_*b*_ = *X*_*i*_ : *y*_*i*_ = *k*, and *k* ∈ *b*} denoting all *n*_*b*_ realized measurements on a given unit (gene) from samples *i* whose condition status *y*_*i*_ maps them to a block *b* of partition *π*,

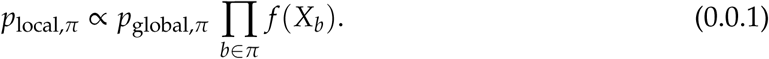

The proportionality is resolved by summing over all *B*_*K*_ partitions, which presents a computational challenge even for moderate *K*. In **EBarrays**, the joint predictive mass *f* (*x*) takes either a compound Gamma form or a log-normal form; our focus is to improve **EBSeq**, which brings empirical Bayes methodology to RNA-Seq data. In **EBSeq**, *f* (*x*) takes a special form as a Beta mixture of Negative Binomial (NB) mass functions. These distributional choices respond to properties and variance characteristics of RNA-Seq data, and also lead to a convenient closed-form predictive mass function:

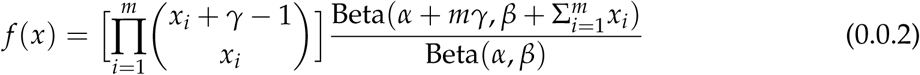

**EBSeq.v1** sets hyperparameters *α, β*, and *γ* as well as mixing rates *p*_global,*π*_ using both local data on the unit and global data on from all units. See Appendix A for additional details. A gene scores highly for pattern *π* if its local posterior probability *p*_local,*π*_ as in (0.0.1) is high, as evaluated through the density (0.0.2). Noting the cancellation of factorial terms, we see that these local scores are determined as a special transform of total expression within each block.

### Pruning

**EBSeq.v1** fits a mixture over all *B*_*K*_ partitions. Even if this were computationally feasible for moderately large *K*, we expect in applications that very many, perhaps most, of these partitions would contribute a negligible amount to the fitted model. We propose a pruning algorithm that uses filtering statistics to identify partitions that are likely to carry most of the mixture mass without the need to fit the mixture over all partitions. The algorithm works by selecting probable partitions at each unit and taking their union as our global pool of partitions possibly of much smaller size than *B*_*K*_.

Consider first a single inference unit (gene) and the group sample means 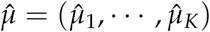:

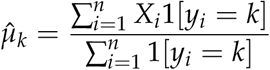

and let *r* = (*r*_1_, …, *r*_*K*_) be the rank of 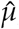, i.e. *r*_*k*_ is the position of 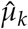 in the permutation rearranging 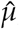 into ascending order. In case of ties we break them according to values of group labels in *y*. We use an overlap concept that was also used in (Dahl 2009) to trim partitions in a modal clustering application.

#### Definition 1.

*Two sets E*_1_ *and E*_2_ *of finite and integer elements overlap if E*_1_ *contains a number between the smallest and largest numbers of E*_2_, *or vice versa.*

For example, *E*_1_ = {1, 3} overlaps with *E*_2_ = {2}, but *E*_1_ = {1, 2} and *E*_2_ = {3} do not overlap. Relative to a partition *π* = {*b*}, consider sets of indices *A*_*r*_(*b*) = {*r*_*k*_, *k* ∈ *b*}.

#### Definition 2.

*Compatibility: A partition π is compatible with an empirical rank r if either π contains only one block or if for any two different blocks b*_1_, *b*_2_ ∈ *π, A*_*r*_(*b*_1_), *A*_*r*_(*b*_2_) *do not overlap.*

For example with *K* = 4, and the empirical ordering 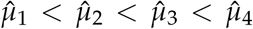 corresponds to rank *r* = (1, 2, 3, 4). The partitions compatible with *r*, and their corresponding hypotheses, are

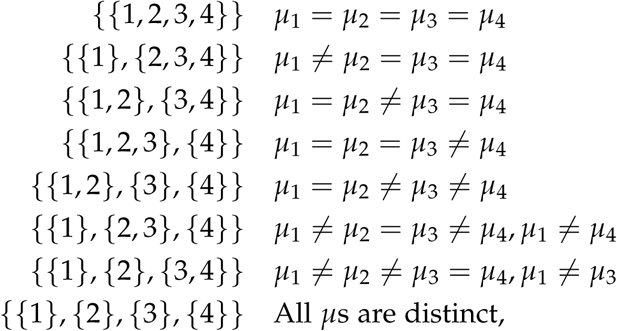

while, for example, *µ*_1_ = *µ*_3_ ≠ *µ*_2_ = *µ*_4_, i.e., {{1, 3}, {2, 4}, is not compatible with *r*. In fact, for the set 𝒞_*g*_ of partitions *π* that are compatible with the empirical ranks at unit *g*, we find:

#### Lemma 1.

*The cardinality of* 𝒞_*g*_ *is* |𝒞_*g*_| = 2^*K*−1^.

Compatible partitions vary from unit to unit, and over the full system there may still be close to *B*_*K*_ different partitions that are compatible for at least one unit. But compatibility is a useful first property to consider as we filter the total number of partitions to a manageable number.

Sampling theory offers one argument in support of considering compatible partitions when the total number of samples *n* is large compared to the number of groups *K*. Then, deviations 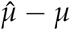 tend to be relatively small. For the pattern *M*_*π*_ corresponding to *µ, π* is not compatible with the empirical ranks only if for some pair of groups *j* and *k*, we have *µ*_*j*_ < *µ*_*k*_ and also 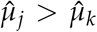. By the law of large numbers, this event is increasingly improbable as *n* increases.

We investigate compatibility empirically by running **EBSeq.v1** in three example data sets where *K* is sufficiently small that computations are feasible: GSE45719 (Deng, Ramsköld, Reinius, and Sandberg 2014), GSE57872 (Patel, Tirosh, Trombetta, Shalek, Gillespie, Wakimoto, Cahill, Nahed, Curry, Martuza *et al.* 2014), GSE74596 (Engel, Seumois, Chavez, Samaniego-Castruita, White, Chawla, Mock, Vijayanand, and Kronenberg 2016). For each data set, Table 1 reports

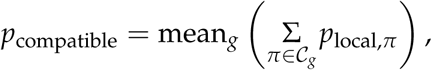

which measures the posterior probability mass of compatible partitions in a fitted model. Compatible partitions cover most of the mixture mass in these cases. Table 1 also reports *N*_95%_, which measures the concentration of local posterior probability mass over the full set of partitions. Specifically, at each unit *g* we consider the most probable partitions and we count how few of these are required to capture at least 95% of the local probability mass, we then average over units to get *N*_95%_. We also keep track of *N*_*c*,95%_, which measures the average number of compatible partitions within the set capturing at least 95% local mass. Notice that *N*_*c*,95%_ and *N*_95%_ are much smaller than |𝒞_*g*_| = 2^*K*−1^, but further pruning is typically necessary to have a manageable number of partitions when considering all units at once.

**Table 1:** Empirical properties of partitions in several data sets. *N*_samples_, *N*_units_ are numbers of samples and units

| Data Set | $N_{\text{samples}}$ | $N_{\text{units}}$ | $K$ | $p_{\text{compatible}}$ | $B_K$ | $N_{95\%}$ | $N_{c,95\%}$ |
| --- | --- | --- | --- | --- | --- | --- | --- |
| GSE45719 | 110 | 27083 | 4 | 95% | 15 | 4.9 | 3.56 |
| GSE74596 | 114 | 23337 | 6 | 82% | 203 | 13.11 | 6.57 |
| GSE52529 | 143 | 22112 | 7 | 85% | 877 | 40.5 | 21.2 |

A key observation is that some pairs of groups present strong information that we can use to filter partitions. For example, at typical levels of variation seen in RNA-Seq data, observing 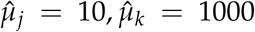 at two groups would suggest *µ*_*j*_ ≠ *µ*_*k*_, and we may not lose much by dropping partitions *π* asserting the opposite: *µ*_*j*_ = *µ*_*k*_. Such information is measured by a Bayes factor comparing differential mean with equivalent mean, namely

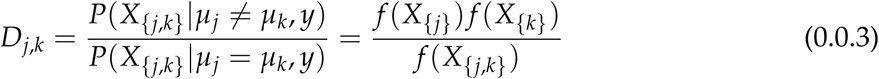

where *X* _{*j*}_ = {*X*_*i*_ : *y*_*i*_ = *j*} denotes all measurements from any group *j, X* _{*j,k*}_ represents vector of all expression values at a given unit for samples from two groups *j* and *k*, and *f* is the predictive density function from (0.0.2). If the two-group Bayes factor is sufficiently extreme (small or large), we are guided about partitions that may be dropped or included before doing a full mixture computation.

In restricting to compatible partitions 𝒞_*g*_, the two-group Bayes factors are relevant to pairs of groups that are adjacent after ranking by empirical mean. Let *o*_*j*_ be the antirank representing the group label having *j*th smallest sample mean. Then each *π* ∈ 𝒞_*g*_ is set by filling either “≠” or “=” into the slots of equation (0.0.4),

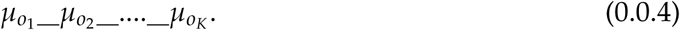

These *K* − 1 slots may be assessed using two-group Bayes factors. To identify a unit specific set of pruned partitions 𝒮_*g*_, we consider three assignment states for each slot in (0.0.4): (equivalent, differential, and uncertain) based on the Bayes factor and a user-specified threshold *t*_1_ > 0.

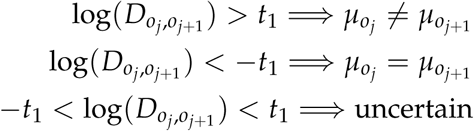

Those partitions consistent with the filled-in (0.0.4) constitute a restricted (i.e., pruned) set of partitions 𝒮_*g*_ ⊂ 𝒞_*g*_. The user-specified threshold *t*_1_ gauges the size of 𝒮_*g*_, with larger *t*_1_ being more inclusive and smaller values being more restrictive. We take 𝒮 = ∪_*g*_𝒮_*g*_ to be the selected set of partitions to be used in the full model-fitting computation. Pseudo-code for the resulting `pruning` tool is in Algorithm 1. For Bayes-factor evaluation, we set hyperparameters (*α, β*) to default values and estimate *γ*; numerical experiments in Appendix C.2 show that 𝒮 is robust to the choice of these hyperparameters.

To investigate `pruning`, we run it on two data sets that are too large for **EBSeq.v1**: Hrvatin (Hrvatin, Hochbaum, Nagy, Cicconet, Robertson, Cheadle, Zilionis, Ratner, Borges-Monroy, Klein *et al.* 2018), Retina (Shekhar, Lapan, Whitney, Tran, Macosko, Kowalczyk, Adiconis, Levin, Nemesh, Goldman *et al.* 2016), and we set the threshold *t*_1_ = 1. We say *N*_UC,*g*_ is the number of uncertain positions at unit *g* and take *N*_selected_ = |𝒮|. Table 2 shows that units *N*_UC,*g*_ is quite small for this threshold setting. Further, we provide users an option to bound the number of uncertainty positions at each unit in order to reduce complexity; details can be found in the Appendix C.2.

**Table 2:** Properties of two-group Bayes factor filtering in two example data sets

| Data Set | $N_{\text{samples}}$ | $N_{\text{units}}$ | $K$ | mean $N_{\text{UC},g}$ | max $N_{\text{UC},g}$ | $N_{\text{selected}}$ | $B_K$ (million) |
| --- | --- | --- | --- | --- | --- | --- | --- |
| Hrvatin | 2164 | 17228 | 12 | 0.016 | 4 | 521 | 4.2 |
| Retina | 15551 | 22156 | 14 | 0.008 | 4 | 376 | 190.9 |

We also check `pruning` in the smaller examples from Table 1. Here we calculate the overall mixture mass (from the full model and **EBSeq.v1**) that is associated with the selected set of partitions: 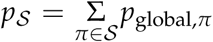. Table 3 confirms that `pruning` finds the dominant mixture components.

**Table 3:**
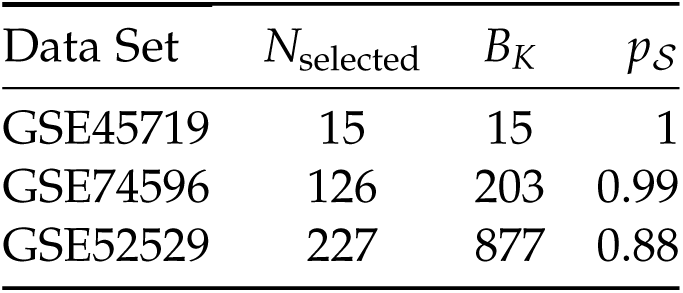
Empirical properties of selected partitions in three example data sets.

Since `pruning` utilizes pairwise comparisons between sample groups in order to select partitions for full model fitting, it is reassuring to know that this pairwise information is sufficient to detect differences that would be found had we accounted for all partitions of all groups. We find:

#### Theorem 1.

*Let hyperparameters α, β, γ be fixed positive integers. If empirical means* 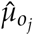 *and* 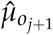 *of adjacent groups o*_*j*_ *and o*_*j*+1_ *are sufficiently distinct, then the partition π*^∗^ *that maximizes P*(*X*|*M*_*π*_, *y*) *over all B*_*K*_ *partitions must place o*_*j*_ *and o*_*j*+1_ *into different blocks.*

Conditions assuring that 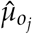 and 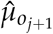 are sufficiently distinct are presented in Appendix B. This finding confirms that the use of pairwise information, such as in `pruning`, conveys properties about the optimal solution one would compute in considering all *B*_*K*_ partitions, without going to the trouble of doing a full survey.

### Crowding

In contrast to `pruning`, we may also seek to prevent scenarios in which too few patterns are selected. This happens when `pruning` creates a chain of equality states even though groups at either end of the chain are significantly different. Then, `pruning` would select

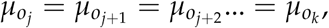

if the two-group Bayes factors favor equality at all adjacent pairs, even though the empirical difference between head and tail of the chain may be large. For example, if 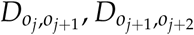 are both near zero and favor equivalent means, then `pruning` selects 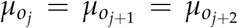. However 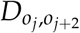 may support uncertain or differential states between these non-adjacent groups. To address this crowding phenomenon, `equalHandle` is an algorithm to break down equality chains when necessary. It adopts an idea from agglomerative hierarchical clustering by iteratively checking the state of adjacent groups with the smallest Bayes factor along the chain. If the state is equal, `equalHandle` combines adjacent groups to form a new, larger group. Otherwise it breaks the chain. Figure 1 demonstrates the algorithm on an equality chain of length 5; see pseudo-code for `equalHandle` in Algorithm 2.

**Figure 1:**
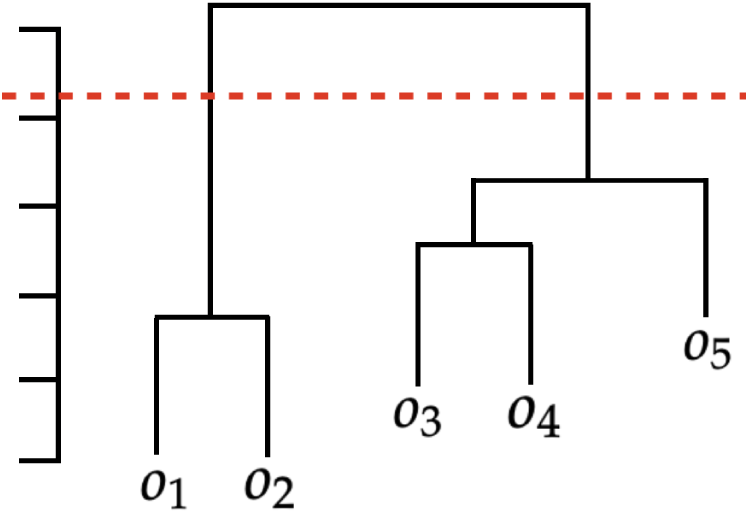
`equalHandle` illustration: *o*_1_, …, *o*_5_ are groups ordered along a chain of equalities suggested by `pruning`. The algorithm sequentially merges adjacent groups having the smallest Bayes factor (strongest evidence for having equal mean) to build a dendrogram. The red line is a threshold where we break the chain, in this case giving two clusters: groups within the same cluster are inferred to have same mean, while the state between clusters remains uncertain, thus adding to the number of possible partitions.

### EBSeq.v2

**EBSeq.v2** works as follows: a pool of partitions is compiled by running `pruning` and `equalHandle` over all units; then the EM algorithm is applied to fit a discrete mixture over this selected pool. Compared to **EBSeq.v1**, we also simplify the optimization within the M-step. **EBSeq.v1** uses `optim` to update the hyperparameters (*α, β*). There is a closed formula for updated mixing rates but hyperparameter optimization by `optim` is more compute intensive. **EBSeq.v2** uses one gradient step toward the optimal solution within each EM iteration. Numerical experiments in Section 3 demonstrate the two methods produce comparable results. Further, **EBSeq.v1** is implemented primarily in R, which, like any other interpreted language, is relatively slow for intensive computing. We deploy core components of **EBSeq.v2** in C++ for efficiency.

#### Algorithm 1 pruning

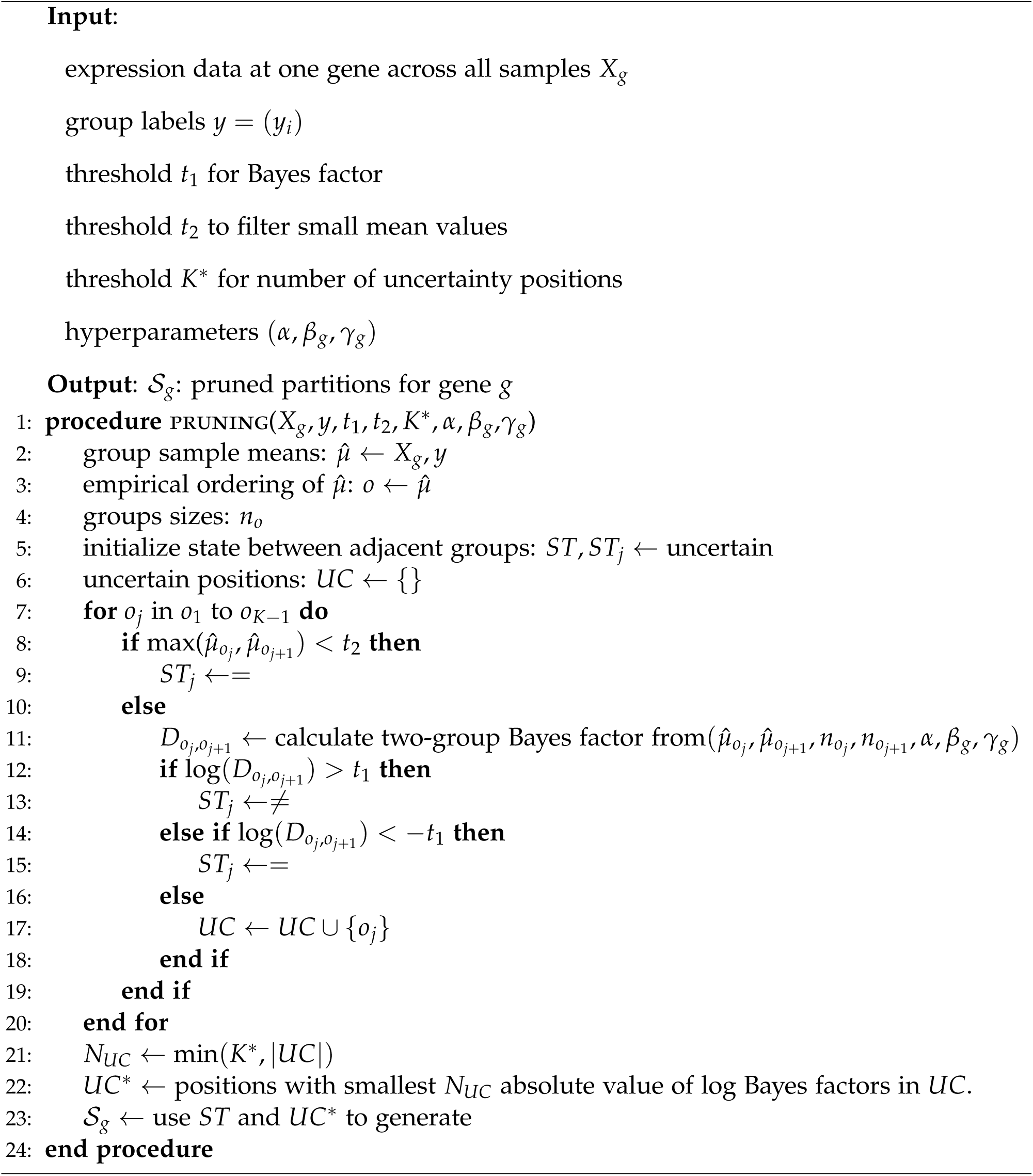

#### Algorithm 2 equal Handle

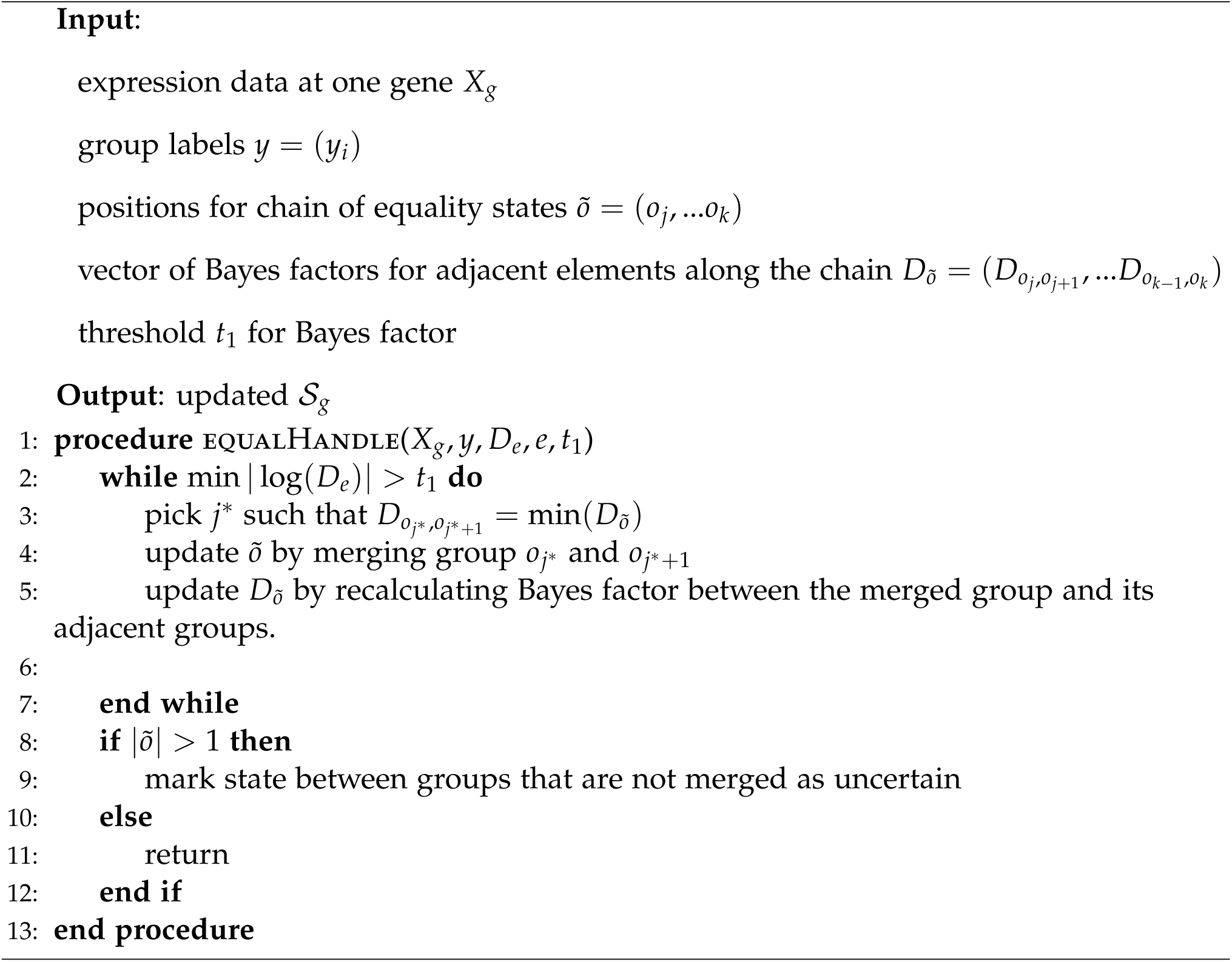

Some applications may require flexibility to further reduce the space of partitions. For example, low intrinsic dimensionality of expression data may effectively reduce the number of relevant partitions (Lopez, Regier, Cole, Jordan, and Yosef 2018). In **EBSeq.v2**, we provide an option for users to prune more aggressively, by iteratively removing partitions having small *p*_global,*π*_. The complete framework is presented as Algorithm 3.

#### Algorithm 3 EBSeq.v2

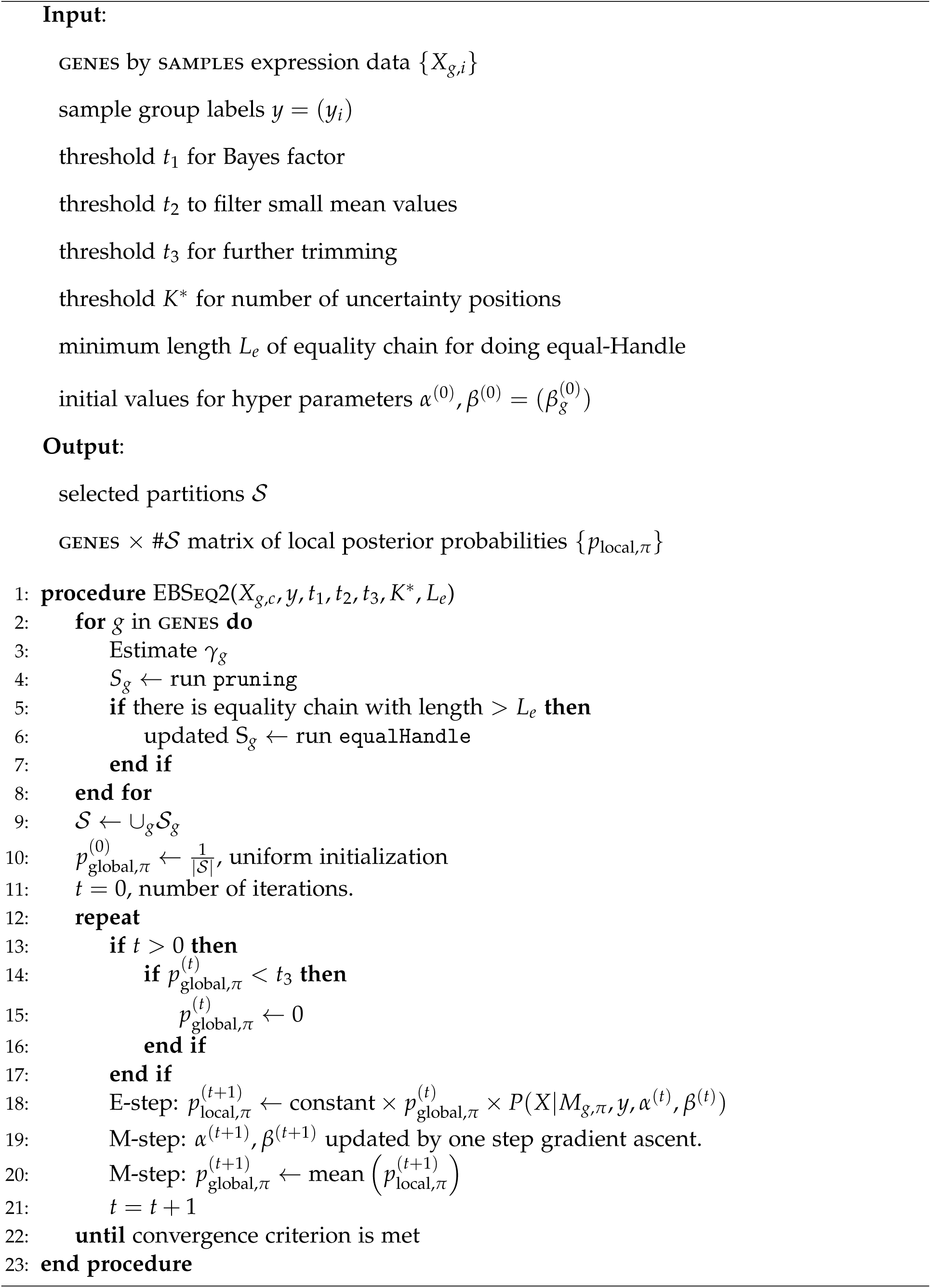

## RESULTS

We compare package versions **EBSeq.v2** and **EBSeq.v1** on benchmark data sets where the number of groups *K* is small. We go on to assess the performance of **EBSeq.v2** in both synthetic and empirical data sets where *K* is larger.

### Benchmarks with small *K*

We check **EBSeq.v2** on the benchmark data sets built into **EBSeq.v1**. Since *K* ≤ 3 in these cases, we use no pruning in deploying **EBSeq.v2**, and so this tests speed and the effect of hyper-parameter optimization differences between the algorithms.

~~~
*R*> *library(EBSeq)*
*R*> *data(“GeneMat”, package = “EBSeq”)*
~~~

This provides a simulated matrix of 1000 rows representing genes and 10 columns representing samples. The columns are ordered such that first and last five columns are corresponding to two different conditions. For both original and new packages, function `EBTest` allows user to get posterior inference of DE patterns among two groups.

The codes for old version

~~~
*R*> *Sizes* <*- MedianNorm(GeneMat)*
*R*> *EBOut* <*- EBTest(Data = GeneMat, Conditions =*
*+ as.factor(rep(c(“C1”,”C2”),each = 5)), sizeFactors = Sizes, maxround = 5)*
*R*> *EBDERes* <*-GetDEResults(EBOut, FDR = 0.05)*
~~~

The codes for new version

~~~
*R*> *Sizes* <*- MedianNorm(GeneMat)*
*R*> *EBOutNew* <*- EBTest(Data = GeneMat, Conditions =*
*+ as.factor(rep(c(“C1”,”C2”),each = 5)), uc = 1*,
*+ sizeFactors = Sizes, maxround = 50*,
*+ step1 = 1e-6, step2 = 1e-2, stopthre = 1e-5)*
*R*> *EBDEResNew* <*- GetDEResults(EBOutNew, FDR = 0.05)*
~~~

The proposed method expects normalized data: we provide a median normalization (Love, Huber, and Anders 2014) in case the input data are not normalized. Basically, the function `MedianNorm` calculates the size factors that are further used for normalization and are passed in the `EBTest` function. **EBSeq.v2** inherits all the functions from version 1, but introduces some extra parameters. For example, we add `uc` to represent the number of uncertainty relation between groups’ means for each gene; `step1` and `step2` are the step-sizes of gradient ascent for optimizing the hyper-parameters (*α, β*) of the beta prior; `maxround` and `stopthre` are parameters to determine the termination of the EM algorithm.

With two groups, the primary outputs of **EBSeq** (either version) are gene-specific local posterior probabilities of equivalent and differential expression – so-called EE and DE probabilities. EE corresponds to the partition *π* having a single block holding all samples (all samples have the same gene-specific expected value). Conditional upon the data, the expected number of EE genes on a list is the sum of the local posterior probabilities (i.e. local false discovery rates); hence we select genes where this EE pattern probability is low.

~~~
*R*> *head(EBDEResNew$PPMat)*
   PPDE PPEE
Gene_1 1 0.0000000e+00
Gene_2 1 7.826157e-47
Gene_3 1 1.162766e-218
Gene_4 1 2.243618e-33
Gene_5 1 3.526946e-229
Gene_6 1 1.250663e-09
~~~

The correlation between the PPDE under two algorithms is quite high: 0.995 Consider a second example:

~~~
*R*> *data(“MultiGeneMat”, package = “EBSeq”)*
~~~

For this dataset, we have 500 genes *n* = 6 samples and *K* = 3 groups (2 samples per group). The codes for old version

~~~
*R*> *MultiSize* <*- MedianNorm(GeneMat)*
*R*> *Conditions* <*- c(“C1”,”C1”,”C2”,”C2”,”C3”,”C3”)*
*R*> *PosParti* <*- GetPatterns(Conditions)*
*R*> *MultiOut* <*- EBMultiTest(Data = MultiGeneMat, NgVector = NULL*,
*+ Conditions = Conditions, AllParti = Parti, sizeFactors = MultiSize, maxround = 5)*
*R*> *MultiPP* <*- GetMultiPP(MultiOut, FDR = 0.05)*
~~~

The codes for new version

~~~
*R*> *MultiSize* <*- MedianNorm(GeneMat)*
*R*> *Conditions* <*- c(“C1”,”C1”,”C2”,”C2”,”C3”,”C3”)*
*R*> *MultiOutNew* <*- EBMultiTest(Data = MultiGeneMat, Conditions =*
*+ Conditions, uc = 2, NgVector = rep(1,nrow(MultiGeneMat))*,
*+ sizeFactors = MultiSize, maxround = 50*,
*+ step2 = 1e-4, stopthre = 1e-5)*
*R*> *MultiPPNew* <*- GetDEResults(MultiOutNew, FDR = 0.05)*
~~~

Function `EBMultiTest` allows user to get posterior inference on DE patterns between multiple groups (more than 3). In the original package, function `GetPatterns` is required for generating all possible patterns that are input in function `EBMultiTest`. In the new package, we get patterns from our pruning algorithm and there is no need to input the set of patterns. At each gene, we consider the patterns *π* with maximum *p*_local,*π*_ under the two algorithms and use adjusted rand index (ARI) to measure their similarity.

The average ARI over genes is 0.961 and close to 1, which indicates that the results from **EBSeq.v1** and **EBSeq.v2** are very similar. We increase the number of simulated genes to 10000 and compare the running time of two algorithms under *K* = 2, 3. Even at small *K*, **EBSeq.v2** is a lot faster than **EBSeq.v1** (Table 4)

**Table 4:** Average run time comparison, 20 samples per group, 10000 genes. EM iteration for **EBSeq.v1** is set to 5

| EBSeq version | $K$ | Average Run Time (minutes) |
| --- | --- | --- |
| v1 | 2 | 3.3 |
| **v2** | **2** | **0.05** |
| v1 | 3 | 45 |
| **v2** | **3** | **0.05** |

**Table 5:**
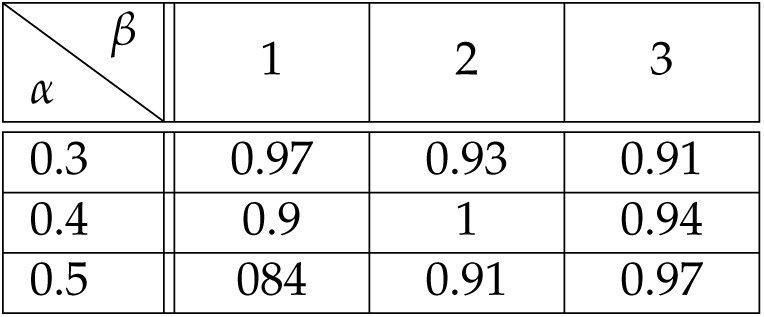
ℐ for comparing 𝒮 under different hyperparamters to the default setting (*α* = 0.4, *β* = 2). Results are averaged over data sets GSE74596 and GSE57872

| $\alpha \backslash \beta$ | 1 | 2 | 3 |
| --- | --- | --- | --- |
| 0.3 | 0.97 | 0.93 | 0.91 |
| 0.4 | 0.9 | 1 | 0.94 |
| 0.5 | 0.84 | 0.91 | 0.97 |

### Synthetic data, larger *K*

We use the Chinese restaurant process (CRP) over both genes and samples to simulate synthetic expression data with plausible variation and clustering characteristics. We report numerical experiments with 20000 genes and 200 samples per group for various numbers of groups *K*; details are in Appendix D.

We evaluate the performance of **EBSeq.v2** using four metrics: coverage, extra patterns, scores and compute time. Let *A* be the set of patterns selected by **EBSeq.v2**, and *B* be the true underlying patterns. `Coverage` is |*A* ∩ *B*|/ |*B*|, `extra patterns` is |*A \ B*|. `Coverage` measures how well we capture the underlying patterns. `Extra patterns` measures the efficiency, namely how many extra patterns we are not able to filter out. For each gene, there is an underlying pattern *π*_*g*_ and an estimated pattern 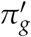 maximizing the posterior (MAP pattern). For the accuracy evaluation, we use the adjusted Rand index (ARI) between *π*_*g*_ and 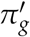. `Time` is the CPU time of running **EBSeq.v2** using a single core 2.4GHz Intel Xeon E5645 with 126 Gb of RAM.

Figure 2 shows operating characteristics for *K* = 15 and *K* = 20 groups (1.38 billion and 51.7 trillion possible patterns). The pruning algorithm covers almost every true, data-generating pattern (Figure 2a). The ARI scores are close to 1 with small standard deviation (Figure 2c,2d), which shows the proposed method can accurately identify generative patterns. The number of extra patterns and compute time are acceptable considering that these cases are beyond the capacity of **EBSeq.v1** (Figure 2b,2e).

**Figure 2:**
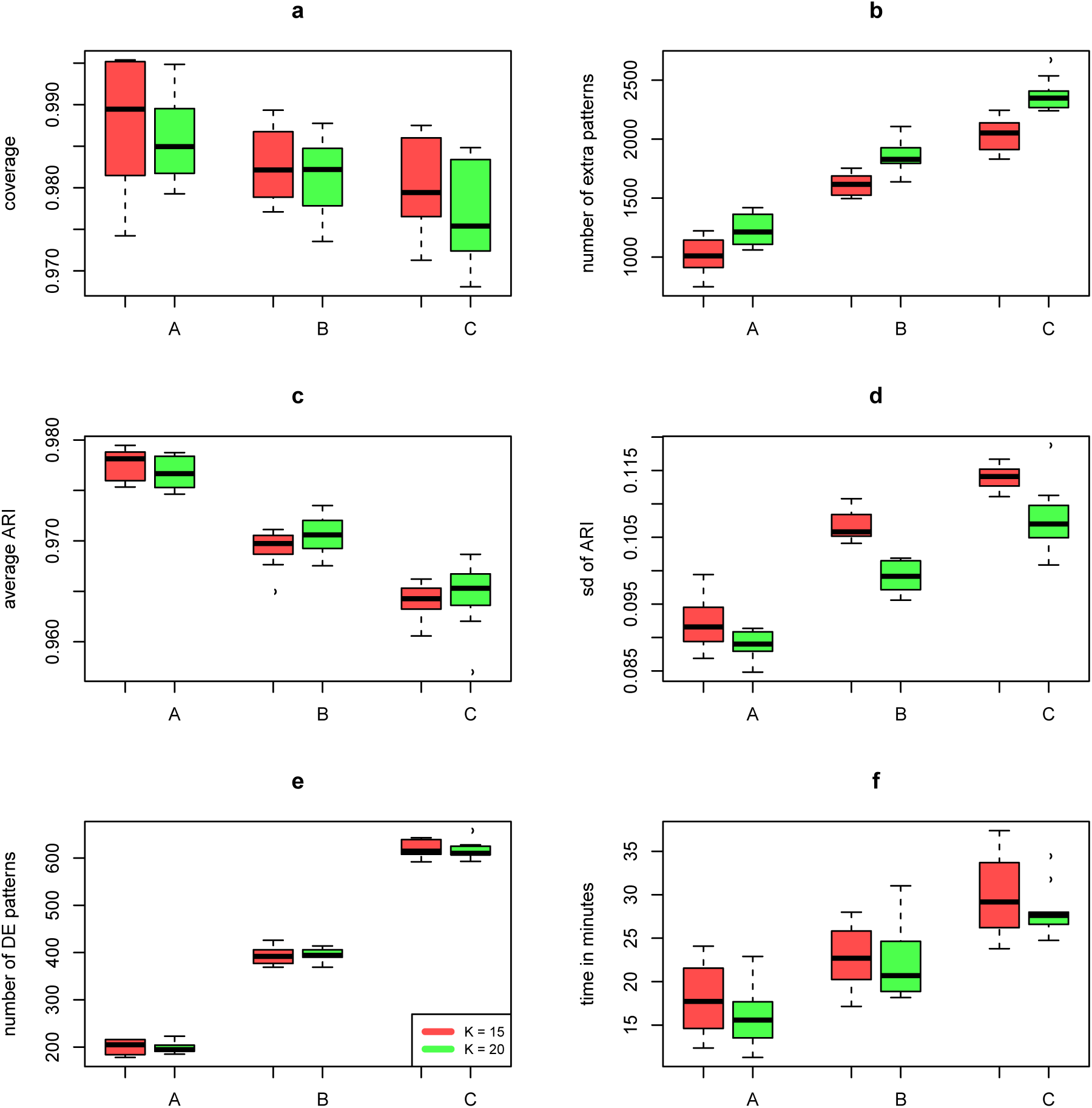
Operating characteristics, synthetic data sets with *K* = 15 (red) or *K* = 20 (green) groups and three settings (horizontal) of a clustering parameter *α*_0_ which affects the expected number of distinct patterns that guide gene-level data. Blocks of genes are generated from Chinese restaurant process, genes within the same block will have same DE patterns. (Approximately 200, 400 and 600 patterns correspond to *A, B*, and *C*, respectively.) Ten synthetic data sets are simulated in each setting, and operating characteristics are computed on each data set. Panel *a* presents the coverage percentage; Panel *b* presents the extra patterns we selected but does not belong to the set of true underlying DE patterns; Panel *c* presents the average ARI (adjusted rand index) between the MAP pattern and true pattern; Panel *d* presents the standard deviation of ARI; Panel *e* presents the number of underlying DE patterns across the genome; Panel *f* presents the computation time (minutes)

### Empirical study

We apply **EBSeq.v2** to three unique-molecule-index (UMI)-based scRNA-seq data sets. These calculations demonstrate the scale of data sets that may be approached with the this new package as well as the sort of inference summaries that become enabled.

The first data set is from a study of mouse visual cortex in which 47, 209 cells were classified into main cell types and subtypes through extensive analysis (Hrvatin *et al.* 2018). We preprocessed the data and applied **EBSeq.v2** to a subset of *n* = 2164 cells and *K* = 12 cell types. Figure 3 shows expression data from a subset of genes, those whose most probable mean expression pattern matches one particular four-block partition. This maximum a posteriori (MAP) pattern *π* is identified by maximizing *p*_local,*π*_ at each gene. As expected by the selection, mean expression (averaging log expression among samples in each group) is similar within block and quite different between blocks of the MAP partition. Sorting genes by their MAP pattern can be a useful filter in applied settings.

**Figure 3:**
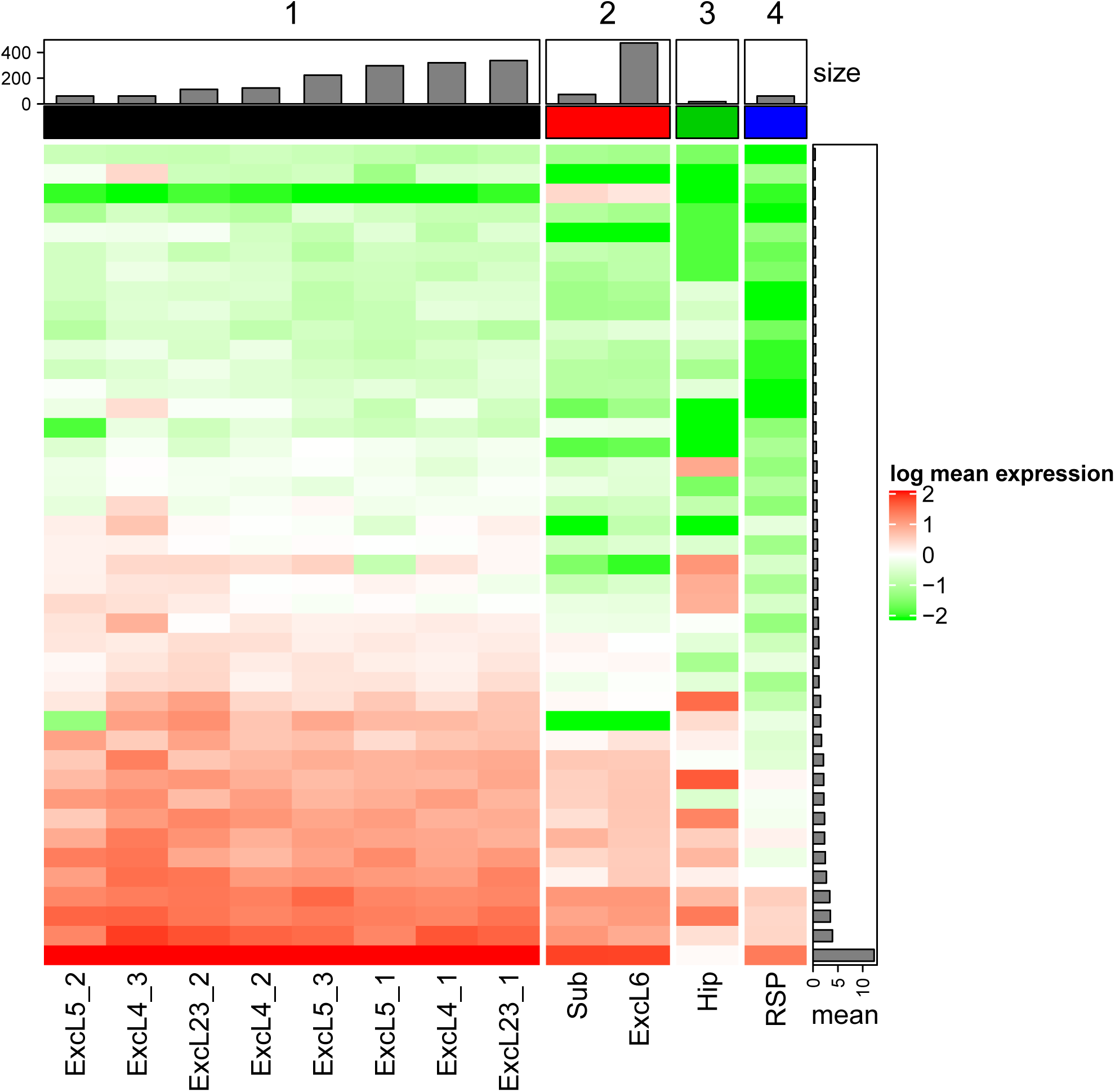
Heatmap of the mean expression on log scale across *K* = 12 groups at those genes with one four-block MAP pattern; genes are filtered by mean expression bigger than 0.4. The top barplot shows the number of cells each group ordered within each block, the right barplot shows the ordered marginal mean across all cells. Data are mouse cortex cells from (Hrvatin *et al.* 2018)

The second data set contains 1000 cells from each of *K* = 10 purified populations derived from peripheral blood^2^. Using **seurat**, (Stuart, Butler, Hoffman, Hafemeister, Papalexi, III, Hao, Stoeckius, Smibert, and Satija 2019), we carried out uniform manifold approximation projection (UMAP) visualizations of the cells considering whole genome (Panel A) versus genes with two particular MAP patterns (Panels B, C) and display results in Figure 4. A single amorphous cluster (Panel B) forms when using genes whose MAP pattern is equivalent expression across all cell types (i.e., the single-block *π* is the MAP pattern for these genes). The UMAP highlights other clusters when we use genes whose MAP pattern matches a partition of interest, such as in Panel C where specific lymphocytes are distinguished. In practice, the use of posterior probabilities for such gene selections can be more sensitive than simpler filters.

**Figure 4:**
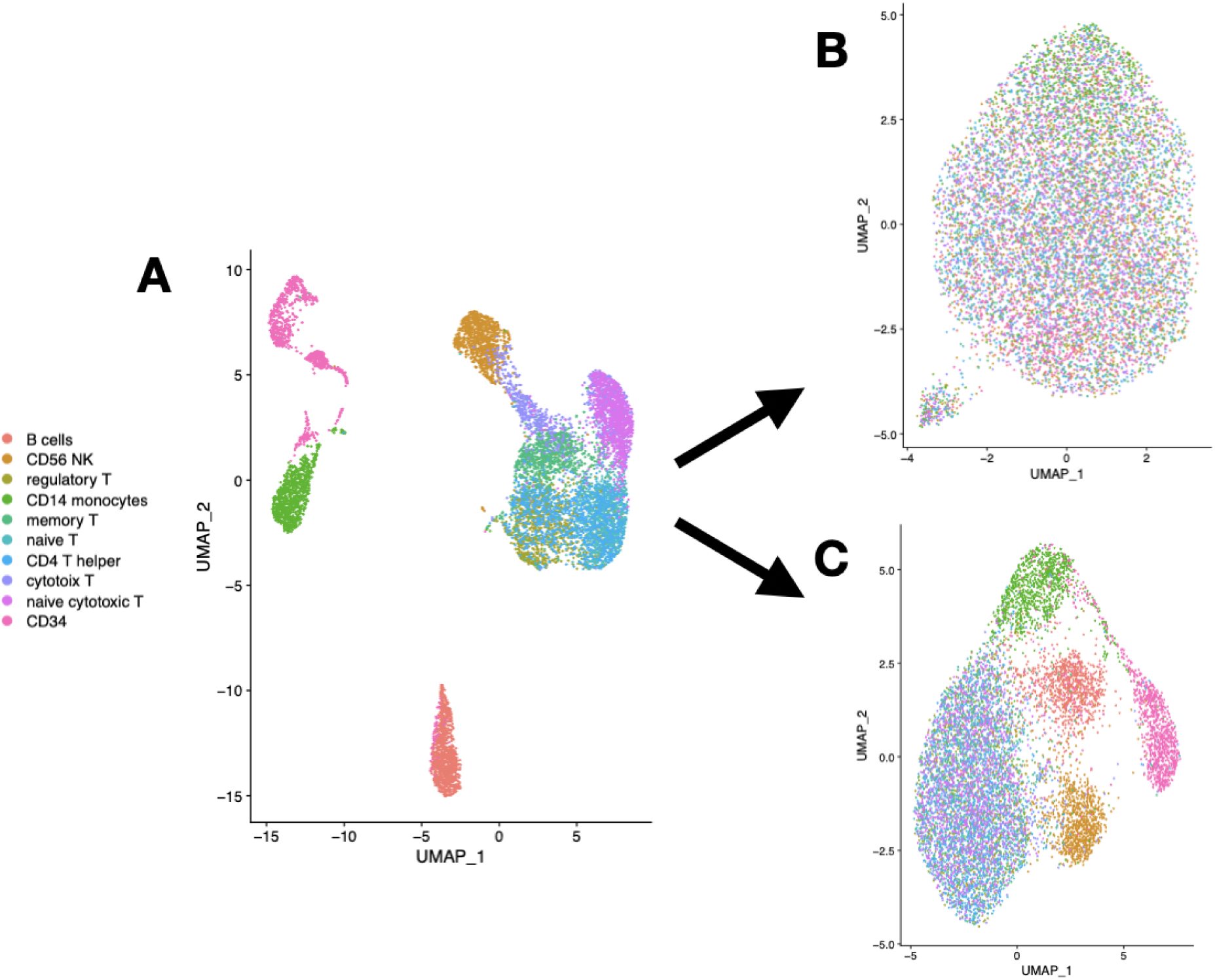
UMAP of PBMC data, left considering whole genome. Panel B only uses genes with null partition as the MAP. Panel C uses genes with MAP patterns distinguishing B cells, CD14, CD34, CD56.

The third data set contained *n* = 27, 499 mouse retinal bipolar cells, we used cluster annotation from *K* = 14 cell types from the author (Shekhar *et al.* 2016). We selected 2 genes “Epha7” and “BC030499” favorable for two partitions and show the cumulative distribution functions (cdfs) across the blocks of these partitions (Figure 5). We observe distributional differences between cells presenting differential means as expected by the selection.

**Figure 5:**
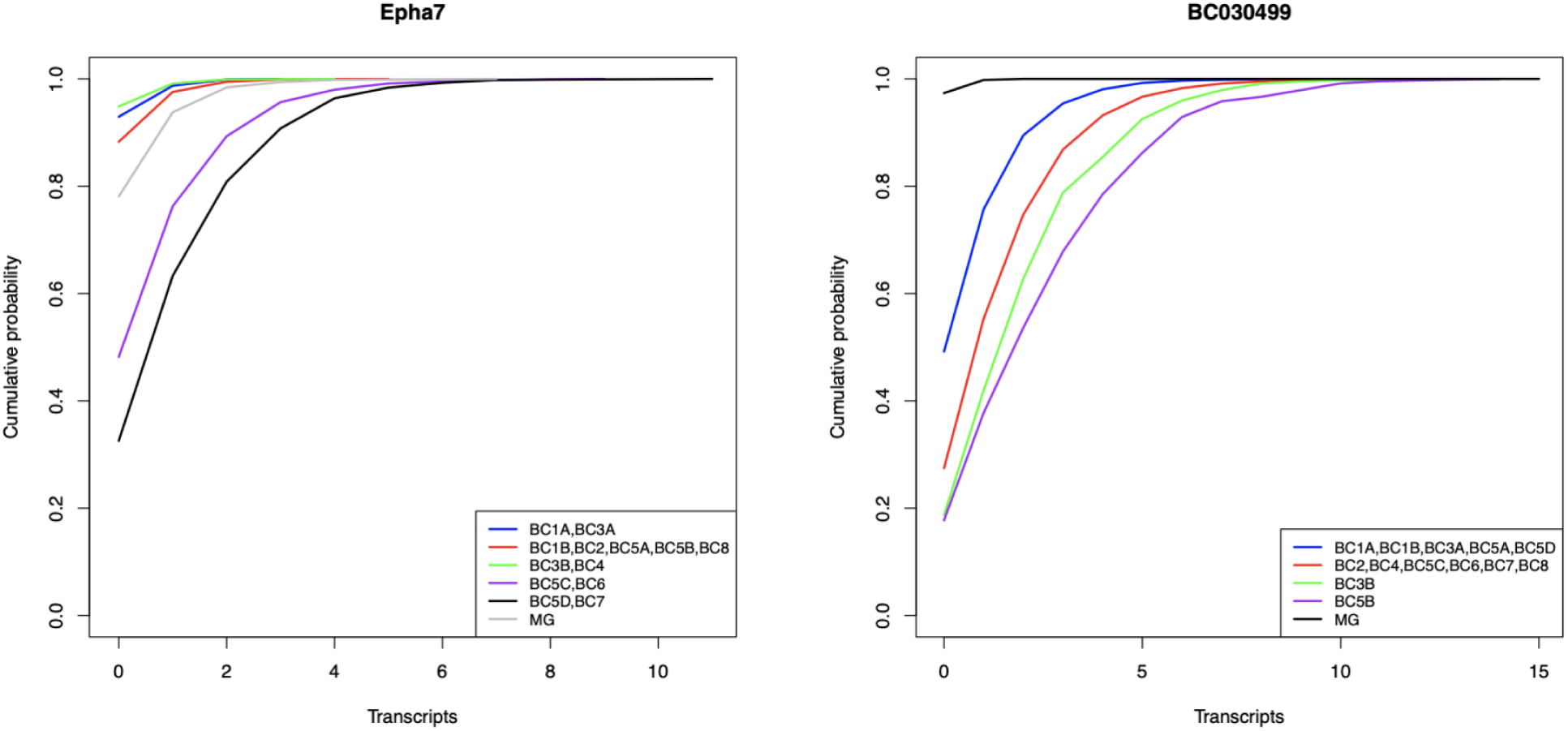
Cumulative distribution of expression at two genes “Epha7” and “BC030499” distinguished by cell condition. Colors represent blocks of conditions sharing the same mean expression in the MAP pattern for these genes. Data are from murine bipolar cells from RETINA (Shekhar *et al.* 2016).

To further check the overall performance of **EBSeq.v2**, we compare pairwise fold change statistics between cellular groups with marginal posterior probabilities of differential expression (DE) between these groups, in the third example with *K* = 12 cellular conditions. Pattern posterior probabilities produced by **EBSeq.v2** are marginalized to get pairwise DE probabilities: 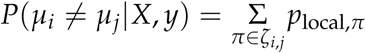 where *ζ*_*i,j*_ is the collection of partitions in which groups *i* and *j* belong to different blocks. Figure 6 compares DE probabilities (below diagonal) to empirical fold changes (above diagonal), here standardized and averaged over the genome, and reveals consistency between these two measures of group difference. Fold changes are often reported, but posterior probabilities have the capacity for self calibration in terms of false discovery rate.

**Figure 6:**
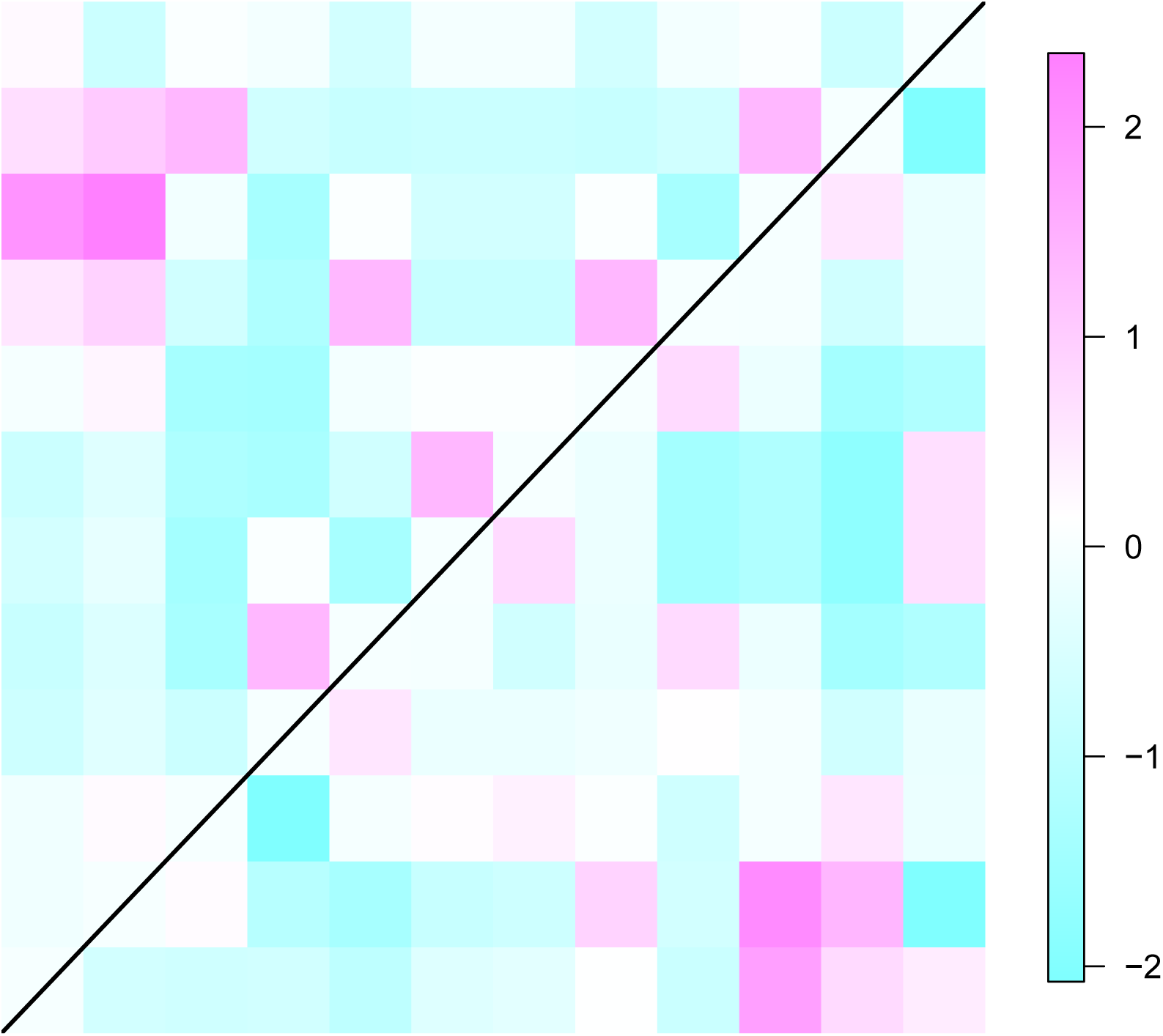
Heatmap shows concordance between standardized fold change (above diagonal) and marginal posterior DE probabilities (below diagonal) between all pairs of *K* = 12 cellular groups using data (Hrvatin *et al.* 2018). Both measures are standardized by centering to zero, scaling to unit standard deviation, and averaging over the genome.

## DISCUSSION

Technologies to measure high-dimensional transcript profiles are still developing, and we expect more large-scale data sets from experiments on immune response, cancer progression, tissue development, and other biological systems. Investigating patterns of gene expression is an essential component of data analysis, e.g., for identifying genes with distributional changes across multiple biological conditions. The mixture of negative binomials model encoded in **EBSeq** approximates the variation of gene expression data in many settings, and the proposed version update extends the domain of application of this important inference tool.

Using an empirical Bayesian approach, **EBSeq** evaluates genes according to likely patterns of differential expression they display over two or more biological conditions. For all genes in a system and for all sorts of patterns of differential expression among sample groups, **EBSeq** scores patterns with local posterior probabilities, and these may be used to measure the false discovery rate of any list of identified genes or to filter genes according to their MAP pattern. In the proposed version 2.0 of **EBSeq**, `pruning` reduces the number of components of the mixture model by virtue of a theoretically guided pairwise Bayes factor calculation, which is key to scaling model-based computations as the number of cellular conditions grows. Also, `equalHandle` deals with a corner case and keeps the model sensitive to the difference between groups. Synthetic and empirical examples demonstrate the efficiency and accuracy of this new version. Future efforts could enable further extensions, such as to allow flexibility in distributional forms while retaining the partition/pattern structure for inference summarization.

## Computational details

The results in this paper were obtained using R 3.5.1 with the packages **Rcpp** 0.12.11, **RcppEigen** 0.3.2.9.0, **BH** 1.69.0-1. R itself and all packages used are available from the Comprehensive R Archive Network (CRAN) at https://CRAN.R-project.org/. **EBSeq.v2** is maintained under the original package name at http://github.com/wiscstatman/EBSeq.

## Acknowledgments

This research was supported in part by NIH grants P50 DE026787, P30CA14520-45, R01 GM102756, NSF grant 1740707, and a Data Science Initiative grant from the University of Wisconsin Office of the Vice Chancellor for Research and Graduate Education. This manuscript was prepared using `overleaf`.

## A. EBSeq margin

A core computation is to evaluate the joint probability mass *f* of input data recordings *X* = (*X*_1_, …, *X*_*m*_) that have a common, unknown, mean, whose value is integrated out (i.e., continuous mixing). In **EBSeq**, we integrate a Negative Binomial mass function, NB(*q, γ*), which has mean *qγ*/(1 − *q*), against a Beta prior for *q*.

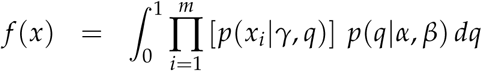

where *x*|*γ, q* is a Negative Binomial(*γ, q*) distribution and *q|α, β* is a Beta distribution with shape parameters *α* and *β*. The result is reported in (0.0.2), where recall Beta(*a, b*) = Γ(*a*)Γ(*b*)/Γ(*a* + *b*), and for *x* > 0, 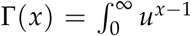 exp{−*u*}*du*. In this representation, the expression data *X* are expected to be normalized and the shape parameter *γ* is common across samples. Thus the conditional mean *µ* is fully determined by *q*.

In **EBSeq** hyperparameters are unit specific, except for *α* which is shared across all units. Different units may have different *β* values, but **EBSeq** allows some blocking of units if they are all isoforms of the same gene, in which case *β* is estimated for the block. Hyper-parameters *γ*_*g*_ are set by the method of moments. The other hyperparameters parameters and the mixing rates *p*_global,*π*_ are estimated via an EM algorithm using data on all samples and all units (Dempster, Laird, and Rubin 1977).

## B. Proofs of Lemma 1 and Theorem 1

### Proof of Lemma 1.

We prove a 1-1 correspondence between compatible partitions and ways of assigning “=” or “≠” in each slot in equation (0.0.4). The cardinality 2^*K*−1^ follows immediately. Without loss of generality, we label groups so that the empirical rank vector *r* is *r* = (1, 2, …, *K*) and the sample means are ordered, 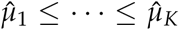. Such labeling induces *A*_*r*_(*b*) = *b*. We fix partitions by resolving slots in equation (0.0.4), which becomes

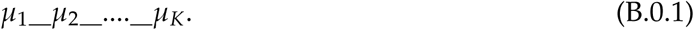

Denote the number of blocks in *π* as *N*(*π*). For the trivial case *N*(*π*) = 1, *π* can be represented by assigning equality “=” to all slots in (B.0.1). If some partition *π* with *N*(*π*) > 1 is compatible with *r*, then *A*_*r*_(*b*_*j*_), *A*_*r*_(*b*_*k*_) are not overlapping for any two blocks *b*_*j*_, *b*_*k*_ ∈ *π*, and so either max(*b*_*j*_) < min(*b*_*k*_) or max(*b*_*k*_) < min(*b*_*j*_). (These extrema are taken over group labels within the block.) For convenience, denote *v*_*j*_ = min(*b*_*j*_) and *w*_*j*_ = max(*b*_*j*_) and let blocks *b*_1_, …, *b*_*N*(*π*)_ of this compatible partition be labeled so they are ordered by the largest element: *w*_1_ ≤ *w*_2_ … ≤*w*_*N*(*π*)_. Since *b*_*j*_ and *b*_*k*_ are not overlapping, we have 1 = *v*_1_ ≤ *w*_1_ < *v*_2_ ≤ *w*_2_ < *v*_*N*(*π*)_ ≤ *w*_*N*(*π*)_ = *K*, and further *v*_*j*+1_ = *w*_*j*_ + 1, *j* = 1, … *N*(*π*) − 1. Thus, the compatible *π* induces the assignment:

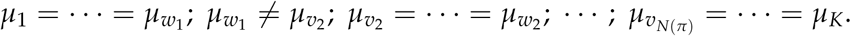

To prove the opposite direction, consider any assignment of “≠” or “=” for the slots in equation (B.0.1), and define partition *π* by the rule that group indices are in the same block if and only if they have equal means by this assignment. If *π* is not compatible with *r*, then it must contain distinct, overlapping blocks *b* and 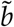 with 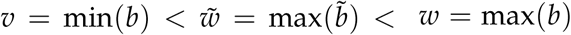. But such an arrangement would imply 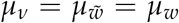, which contradicts the presumption that *w* and 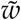 are in different blocks. Thus *π* is compatible with *r*.

□

Without loss of generality, we label adjacent groups by *j* = 1, 2 and recall the notation from Section 2.2 where *X* _{*j*}_ denotes the vector of *n*_*j*_ sample measurements in group *j* at the unit on test. In preparing to prove Theorem 1, we decompose predictive functions in (0.0.2) which are specific mixtures of Negative Binomial masses:

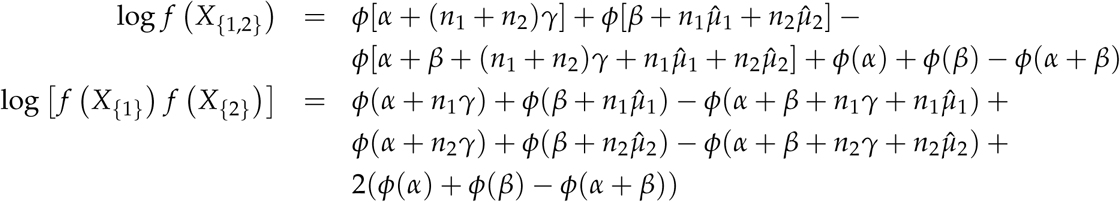

where *ϕ*(*x*) = log Γ(*x*), Γ(*x*) is the usual Gamma function, hyperparameters *α, β*, and *γ* are fixed positive integers, and 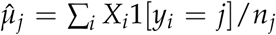 with ordering 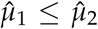. We use this decomposition to understand the two-group Bayes factor (0.0.3), which in this setting is:

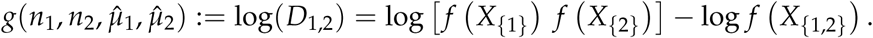

In the notation we suppress dependence of the two-group Bayes factor on the hyperparameters; these are fixed throughout and our analysis rests on changes in *g* with respect to changes in the other inputs. (We also note that in applications the hyperparameters may take positive, non-integer values; we restrict to integers in the theorem only because the proof method is more straightforward.)

### Lemma 2.

*If* 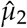 *is sufficiently larger than* 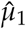, *then g*(*n*_1_,*n*_2_, 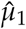, 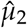) > 0.

### Proof of Lemma 2.

For notation, let 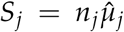, *R*_*j*_ = *n*_*j*_*γ*, for *j* = 1, 2, and recall that for positive integers *x, ϕ*(*x* + 1) = log(*x*!). Manipulating the two-group Bayes factor we obtain:

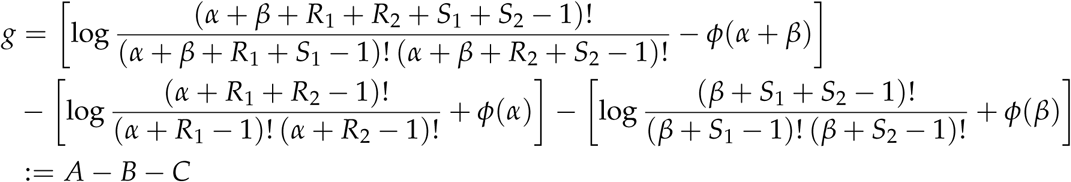

Further we have

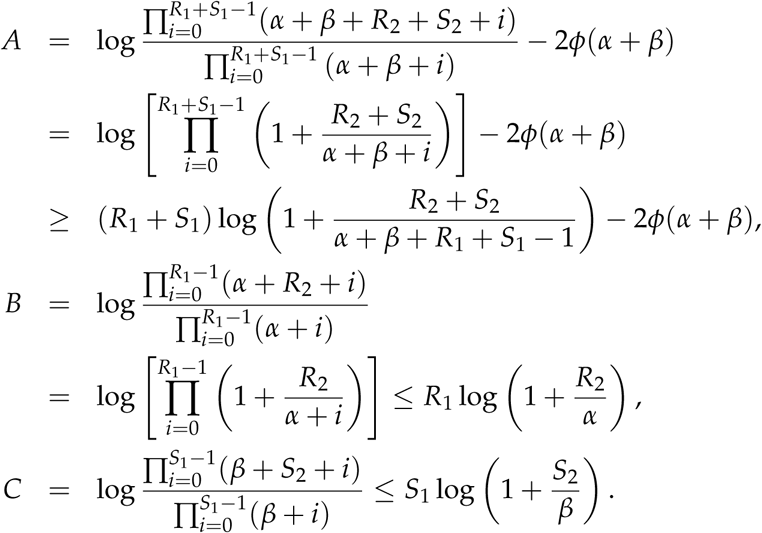

Thus,

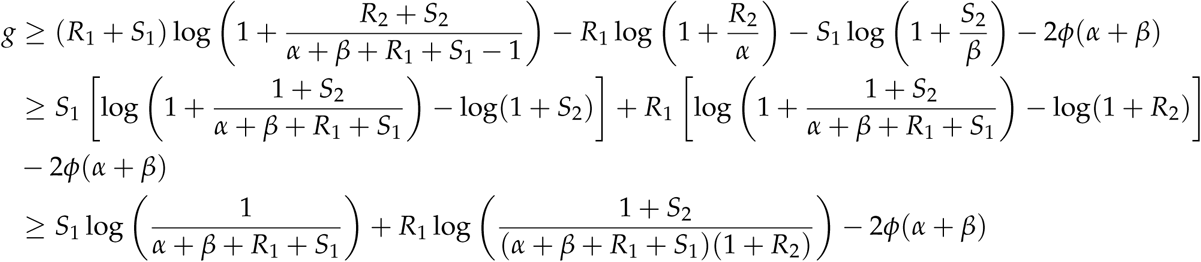

This becomes positive as 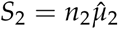 increases.

Recall the digamma function *ψ*(*x*) = *ϕ*′(*x*), and the recursive property of *ψ*:

### Lemma 3.

*For positive real x and natural N, changes in the digamma function ψ satisfy:*

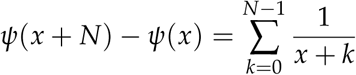

### Lemma 4.

*If* 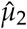 *is sufficiently larger than* 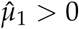, *then g is:*

a. *monotone decreasing in* 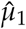
b. *monotone increasing in* 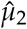
c. *monotone increasing in n*_1_
d. *monotone increasing in n*_2_

*These monotonicities are in one variable at a time with the others fixed.*

### Proof of Lemma 4.

We use notation for *S*_*j*_ and *R*_*j*_ as in the proof of Lemma 2, but treat *g* as a differentiable function of its arguments and study its partial derivatives. To prove a), we have the derivative

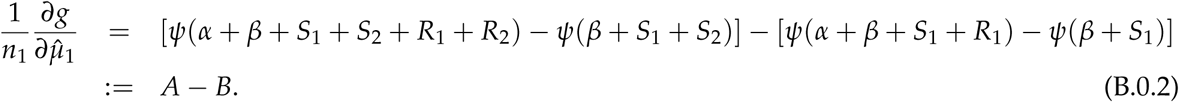

Using the Lemma 3 separately on *A* and *B* leads to

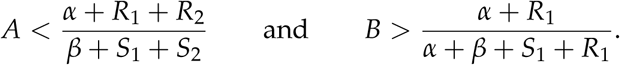

The derivative (B.0.2) is negative if *A* < *B*, which is assured when 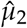 is sufficiently in excess of 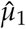 because then 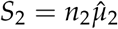 is large enough to push *A* below *B*.

To prove b), the derivative for 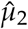 is

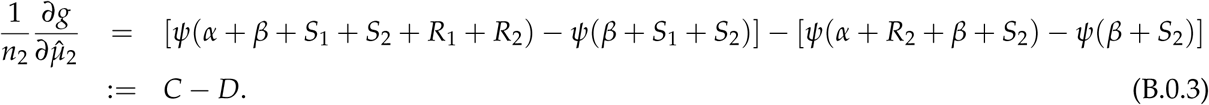

Using Lemma 3 again, we have

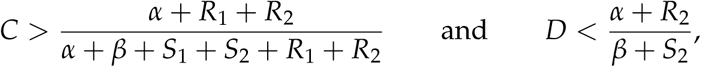

which guarantees positive derivative B.0.3 if

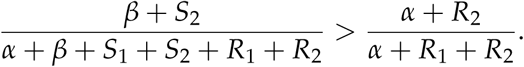

This holds when 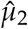 is sufficiently larger than 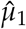 b ecause t he l eft h and s ide a pproaches 1 while right hand side remains bounded below 1. To prove c), the derivative for *n*_1_ is

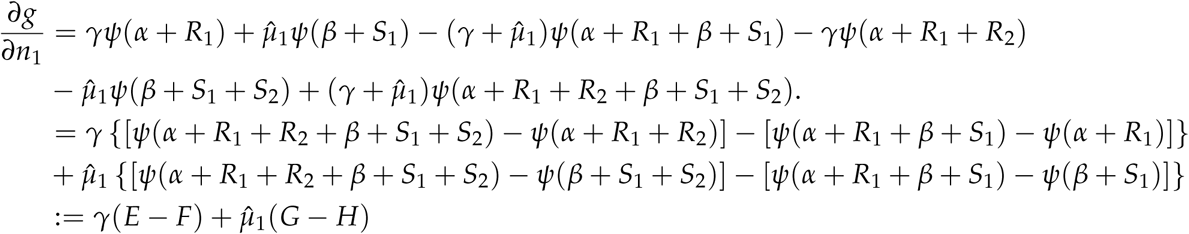

Following the same approach using Lemma 3, we have

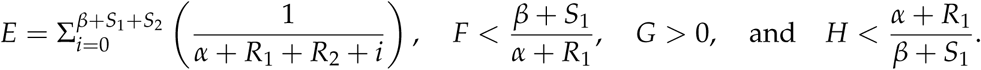

*E* > *F* + *H* is assured for sufficiently large 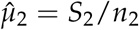 since *E* tracks a divergent sum. Proof of *d*) is similar and details are omitted. □

### Proof of Theorem 1.

Without loss of generality we label the groups 1, 2, …, *K*, so that their empirical means are non-decreasing: 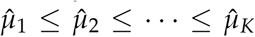. Let *ζ* be the set of all possible partitions of *K* elements (numbering *B*_*K*_), and for some *j* < *K* where 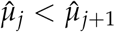, and let *ζ*_*e*_ be the subset of these partitions assigning equivalent means *µ*_*j*_ = *µ*_*j*+1_ to groups *j* and *j* + 1. In other words, every partition in *ζ*_*e*_ is such that *j* and *j* + 1 are elements of the same partition block. Note that sample sizes *n*_*j*_ and *n*_*j*+1_ are associated with these two groups of samples. Further, let *ζ*_*d*_ = *ζ \ ζ*_*e*_ be set of partitions assigning differential means to *j* and *j* + 1. As for partitions in *ζ*_*e*_, we can merge *X*_{*j*}_ and *X*_{*j*+1}_ into one group and view *ζ*_*e*_ as partitions on *K* − 1 elements, so |*ζ*_*e*_| = *B*_*K*−1_, while |*ζ*_*d*_| = *B*_*K*_ *B*_*K*−1_. Therefore, |*ζ*_*e*_| < |*ζ*_*d*_|. There is an injection mapping from *ζ*_*e*_ to *ζ*_*d*_. Any block *b* ∈ *π* ∈ *ζ*_*e*_ that excludes *j* and *j* + 1 is part of the partition *π*′ ∈ *ζ*_*d*_ uniquely associated with *π*. The single block 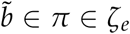 that contains both *j* and *j* + 1 is split into two blocks 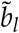 and 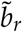 in *π*′; this is the one and only split for which 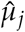 is the largest sample mean in 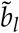 and 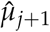 is the smallest sample mean in block 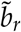; let us say that these two blocks contain *n*_*l*_ and *n*_*r*_ samples, respectively.

In searching for a partition to maximize the the joint predictive mass of all data *X* on the unit in focus, consider the ratio of predictive masses comparing partitions *π* and *π*′ defined above:

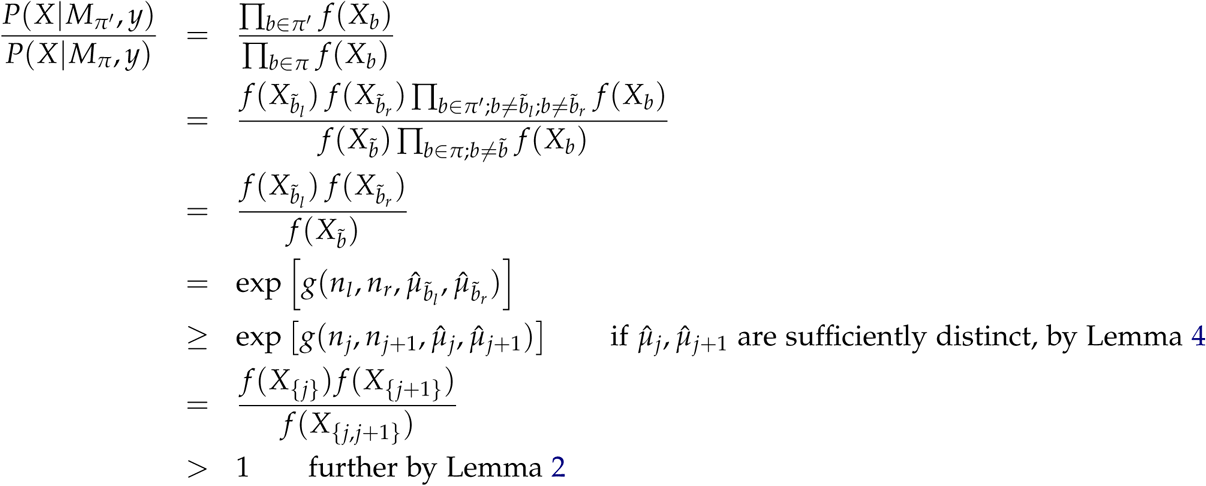

The first inequality above uses the monotonicity in Lemma 4 together with the readily confirmed facts that: 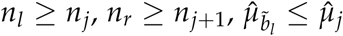, and 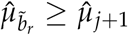.

Thus, the partition with maximal *P*(*X*|*M*_*π*_, *y*) must assign differential means between groups *j* and *j* + 1 when 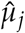 and 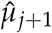 are sufficiently distinct.

□

## C. Further computational details

### C.1. Number of groups *K*

For all the empirical examples used in Section 2, their group number *K* is determined via clustering analysis in **scDDboost** (Ma *et al.* 2019).

### C.2. Hyperparameters of two-group Bayes factor

For different hyperparameters the selected partitions 𝒮 may vary. To measure the similarity between two sets of partitions 𝒮_1_ and 𝒮_2_, we have the metric:

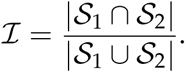

The default value of *α* is 0.4, and *β* is set to 2 for all units.

Table shows that the final selected partitions are robust to the choice of (*α, β*) used in two-group Bayes factor. The scale of *α* and *β* are determined by estimations from **EBSeq.v1** applied on benchmark data sets built into it and an empirical data GSE45719. Similar calculations (not shown) confirm that the two-group Bayes factor is quite insensitive to plausible values of hyperparameter *γ*.

### C.3. Number of uncertain positions

We provide an option to limit the number of uncertain positions. Users can specify a upper bound *U*^∗^ for *N*_UC,*g*_, which is shared by all the units. Let 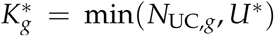. Then for unit *g*, we keep the smallest 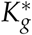 uncertain positions in terms of the absolute value of two-group Bayes factor in the log scale.

## D. Simulation details

In the simulation study we set 200 samples per group and 20000 units (genes). It is biologically plausible that a DE pattern can be shared by several units, so we assign blocks to units and units belong to the same block have same pattern. The blocks are generated from a Chinese restaurant process (CRP) with strength and discount parameter (*α*_0_, *α*_1_) (Aldous, Ibragimov, and Jacod 2006). Namely

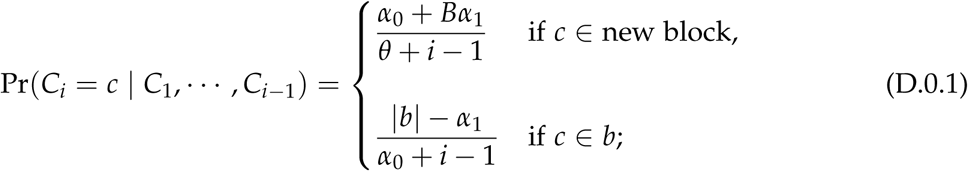

where *C*_1_, …, *C*_*i*_ are the cluster label for elements 1, …, *i. B* is the number of blocks, |*b*| is the number of elements inside that block. We set *α*_1_ = 1 and use *α*_0_ to control the number of blocks. After we obtain the blocking structure of genes, the next step is to generate partitions of groups and assign them to those blocks of genes. One challenge is that the size of possible partitions increases rapidly, for example when *K* = 15, there is 1.38 billion partitions, makes it difficult to sample directly. Again, we use CRP to generate partitions. The remaining problem is to assign partitions to blocks of genes. We observe that in the expression data, partitions *π* with big *p*_local,*π*_ for most genes have small number of blocks and thus are coarse. Therefore, we use a monotone mapping, for example, big block of units will be associated with coarse partitions and small block of units will get fine partitions. We use entropy to measure the complexity(fineness) of a partition. That is, for a partition *π*, we have the entropy: 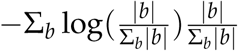, it reaches minimum when all samples belong to a single block and reaches maximum when every sample forms a distinct block. We assign the patterns with low entropy to big blocks. At each unit, counts are sampled from a Beta mixture of Negative Binomial (NB) distributions.

## Footnotes

1 This manuscript version is a technical report from the University of Wisconsin Department of Biostatistics and Medical Informatics, and was posted to bioRxiv July 2, 2020.

2 Available https://support.10xgenomics.com/single-cell-gene-expression/datasets

## References

Aldous DJ, Ibragimov IA, Jacod J (2006). Ecole d’Ete de Probabilites de Saint-Flour XIII, 1983, volume 1117. Springer. ISBN 3540393161.

Baños-Lara MDR, Zabaleta J, Garai J, Baddoo M, Guerrero-Plata A (2018). “Comparative analysis of miRNA profile in human dendritic cells infected with respiratory syncytial virus and human metapneumovirus.” BMC research notes, 11(1), 432.

Boost (2015). “Boost C++ Libraries.” http://www.boost.org/. Last accessed 2015-06-30.

Dahl DB (2009). “Modal clustering in a class of product partition models.” Bayesian Anal., 4(2), 243–264. doi:10.1214/09-BA409. URL https://doi.org/10.1214/09-BA409.

Dempster AP, Laird NM, Rubin DB (1977). “Maximum likelihood from incomplete data via the EM algorithm.” Journal of the Royal Statistical Society: Series B (Methodological), 39(1), 1–22.

Deng Q, Ramsköld D, Reinius B, Sandberg R (2014). “Single-Cell RNA-Seq Reveals Dynamic, Random Monoallelic Gene Expression in Mammalian Cells.” Science, 343(6167), 193–196. ISSN 0036-8075. doi:10.1126/science.1245316. https://science.sciencemag.org/content/343/6167/193.full.pdf, URL https://science.sciencemag.org/content/343/6167/193.

Efron B (2010). Large-Scale Inference: Empirical Bayes Methods for Estimation, Testing, and Prediction. Institute of Mathematical Statistics Monographs. Cambridge University Press. doi:10.1017/CBO9780511761362.

Engel I, Seumois G, Chavez L, Samaniego-Castruita D, White B, Chawla A, Mock D, Vijayanand P, Kronenberg M (2016). “Innate-like functions of natural killer T cell subsets result from highly divergent gene programs.” Nature immunology, 17(6), 728–739.

Gardner M (1978). “Mathematical Games.” Scientific American, 238(5), 24–30. doi:10.1038/scientificamerican0578-24.

Guennebaud G, Jacob B, et al. (2010). “Eigen v3.” http://eigen.tuxfamily.org.

Heller KA, Ghahramani Z (2005). “Bayesian hierarchical clustering.” In Proceedings of the 22nd international conference on Machine learning, pp. 297–304.

Hrvatin S, Hochbaum DR, Nagy MA, Cicconet M, Robertson K, Cheadle L, Zilionis R, Ratner A, Borges-Monroy R, Klein AM, et al. (2018). “Single-cell analysis of experience-dependent transcriptomic states in the mouse visual cortex.” Nature neuroscience, 21(1), 120–129.

Huber W, Carey VJ, Gentleman R, Anders S, Carlson M, Carvalho BS, Bravo HC, Davis S, Gatto L, Girke T, Gottardo R, Hahne F, Hansen KD, Irizarry RA, Lawrence M, Love MI, MacDonald J, Obenchain V, Ole’s AK, Pag’es H, Reyes A, Shannon P, Smyth GK, Tenenbaum D, Waldron L, Morgan M (2015). “Orchestrating high-throughput genomic analysis with Bioconductor.” Nature Methods, 12(2), 115–121. URL http://www.nature.com/nmeth/journal/v12/n2/full/nmeth.3252.html.

Kendziorski C, Newton M, Lan H, Gould M (2003). “On parametric empirical Bayes methods for comparing multiple groups using replicated gene expression profiles.” Statistics in medicine, 22(24), 3899–3914.

Lee CJ, Ahn H, Jeong D, Pak M, Moon JH, Kim S (2020). “Impact of mutations in DNA methylation modification genes on genome-wide methylation landscapes and down-stream gene activations in pan-cancer.” BMC Medical Genomics, 13(3), 1–14.

Leng N, Dawson JA, Thomson James Aand Ruotti V, Rissman AI, Smits BMG, Haag JD, Gould MN, Stewart RM, Kendziorski C (2013). “EBSeq: an empirical Bayes hierarchical model for inference in RNA-seq experiments.” Bioinformatics, 29(8), 1035–1043. ISSN 1367-4803. doi:10.1093/bioinformatics/btt087. https://academic.oup.com/bioinformatics/article-pdf/29/8/1035/17106263/btt087.pdf URL https://doi.org/10.1093/bioinformatics/btt087.

Leng N, Kendziorski C (2019). EBSeq: An R package for gene and isoform differential expression analysis of RNA-seq data. R package version 1.24.0.

Lopez R, Regier J, Cole MB, Jordan MI, Yosef N (2018). “Deep generative modeling for single-cell transcriptomics.” Nature Methods, 15(12), 1053–1058. doi:10.1038/s41592-018-0229-2. URL https://doi.org/10.1038/s41592-018-0229-2.

Louro B, Marques JP, Manchado M, Power DM, Campinho MA (2020). “Sole head transcrip-tomics reveals a coordinated developmental program during metamorphosis.” Genomics, 112(1), 592–602.

Love MI, Huber W, Anders S (2014). “Moderated estimation of fold change and dispersion for RNA-seq data with DESeq2.” Genome biology, 15(12), 550.

Ma BX, Korthauer K, Kendziorski C, Newton MA (2019). “A Compositional Model to Assess Expression Changes from Single-Cell Rna-Seq Data.” bioRxiv. doi:10.1101/655795. https://www.biorxiv.org/content/early/2019/05/31/655795.full.pdf, URL https://www.biorxiv.org/content/early/2019/05/31/655795.

MacEachern SN (1998). “Computational methods for mixture of Dirichlet process models.” In Practical nonparametric and semiparametric Bayesian statistics, pp. 23–43. Springer.

Newhouse DJ, Hofmeister EK, Balakrishnan CN (2017). “Transcriptional response to West Nile virus infection in the zebra finch (Taeniopygia guttata).” Royal Society open science, 4(6), 170296.

O’Grady T, Baddoo M, Flemington EK (2017). “Analysis of EBV Transcription Using High-Throughput RNA Sequencing.” In Epstein Barr Virus, pp. 105–121. Springer.

Patel AP, Tirosh I, Trombetta JJ, Shalek AK, Gillespie SM, Wakimoto H, Cahill DP, Nahed BV, Curry WT, Martuza RL, et al. (2014). “Single-cell RNA-seq highlights intratumoral heterogeneity in primary glioblastoma.” Science, 344(6190), 1396–1401.

Quintana FA, Iglesias PL (2003). “Bayesian clustering and product partition models.” Journal of the Royal Statistical Society: Series B (Statistical Methodology), 65(2), 557–574.

Sabbagh MF, Heng JS, Luo C, Castanon RG, Nery JR, Rattner A, Goff LA, Ecker JR, Nathans J (2018). “Transcriptional and epigenomic landscapes of CNS and non-CNS vascular endothelial cells.” Elife, 7, e36187.

Sanders SM, Cartwright P (2015). “Patterns of Wnt signaling in the life cycle of Podocoryna carnea and its implications for medusae evolution in Hydrozoa (Cnidaria).” Evolution & development, 17(6), 325–336.

Shekhar K, Lapan SW, Whitney IE, Tran NM, Macosko EZ, Kowalczyk M, Adiconis X, Levin JZ, Nemesh J, Goldman M, et al. (2016). “Comprehensive classification of retinal bipolar neurons by single-cell transcriptomics.” Cell, 166(5), 1308–1323.

Son JC, Jeong HO, Park D, No SG, Lee EK, Lee J, Chung HY (2017). “miR-10a and miR-204 as a potential prognostic indicator in low-grade gliomas.” Cancer informatics, 16, 1176935117702878.

Song X, Tang T, Li C, Liu X, Zhou L (2018). “CBX8 and CD96 are important prognostic biomarkers of colorectal cancer.” Medical science monitor: international medical journal of experimental and clinical research, 24, 7820.

Stuart T, Butler A, Hoffman P, Hafemeister C, Papalexi E, III WMM, Hao Y, Stoeckius M, Smibert P, Satija R (2019). “Comprehensive Integration of Single-Cell Data.” Cell, 177, 1888–1902. doi:10.1016/j.cell.2019.05.031. URL https://doi.org/10.1016/j.cell.2019.05.031.

Yang F, Lv SX, Lv L, Liu YH, Dong SY, Yao ZH, Dai Xx, Zhang XH, Wang OC (2016). “Identification of lncRNA FAM83H-AS1 as a novel prognostic marker in luminal subtype breast cancer.” OncoTargets and therapy, 9, 7039.

Yoon KJ, Ringeling FR, Vissers C, Jacob F, Pokrass M, Jimenez-Cyrus D, Su Y, Kim NS, Zhu Y, Zheng L, et al. (2017). “Temporal control of mammalian cortical neurogenesis by m6A methylation.” Cell, 171(4), 877–889.

Yuan M, Newton M, Sarkar D, Kendziorski C (2019). EBarrays: Unified Approach for Simul-taneous Gene Clustering and Differential Expression Identification. R package version 2.50.0.

Zhang Q, Zeng LP, Zhou P, Irving AT, Li S, Shi ZL, Wang LF (2017). “IFNAR2-dependent gene expression profile induced by IFN-α in Pteropus alecto bat cells and impact of IFNAR2 knockout on virus infection.” PloS one, 12(8).

